# On the spillover effect and optimal size of marine reserves for sustainable fishing yields

**DOI:** 10.1101/853879

**Authors:** Nao Takashina

## Abstract

Marine reserves are an essential component of modern fishery management. Marine reserves, which represent a management tradeoff between harvesting and conservation, are fundamental to maintenance of fisheries. Finding optimal reserve sizes that improve fishing yields is not only of theoretical interest, but also of practical importance to facilitate decision making. Also, since the migratory behavior of some species influences the spillover effect of a marine reserve, this is a key consideration when assessing performance of marine reserves. The relationship between optimal reserve size and migration rate/mode has not been well studied, but it is fundamental to management success. Here, I investigate optimal reserve size and its management outcome with different levels of spillover via a simple two-patch mathematical model. In this model, one patch is open to fishing, and the other is closed. The two-patch model is aggregated by single-population dynamics when the migration rate is sufficiently larger than the growth rate of a target species. At this limit, I show that an optimal reserve size exists when pre-reserve fishing occurs at fishing mortality larger than *f*_*MSY*_, the fishing mortality at the maximum sustainable yield (MSY). Also, the fishing yield at an optimal reserve size becomes as large as MSY at the limit. Numerical simulations at various migration rates between the two patches suggest that the maximum harvest under management with a marine reserve is achieved at this limit. This contrasts with the conservation benefit which is maximized at an intermediate migration rate. Numerical simulations show that the above-mentioned condition derived from the aggregated model is necessary when the migration rate is not sufficiently large, and that a moderate migration rate is further necessary for an optimal reserve size to exist. However, high fishing mortality reduces this requirement.

## 1 Introduction

Marine reserves, or no-take marine protected areas (MPAs) are a central tool in modern fishery management to reduce fishing pressure and to preserve biodiversity [27, 46, 56]. Implementation of marine reserves leads to fishing closures, and it can cause a management tradeoff between harvest and conservation [10, 33]. The spillover effect from marine reserves is essential to characterize fishery and conservation benefits of reserves [22]; hence, understanding the migration effect is fundamental to management success.

In modern fishery management, in which managers seek optimal yields [11, 30], optimal reserve sizes are widely discussed [16, 25, 39]. Optimal yield is also a theoretically important concept, as it represents a baseline to assess the intensity of a management tradeoff. Marine reserves can reduce fishing yields when a fishery manages a stock sustainably [25, 28, 42, 59]; hence, seeking an optimal reserve size is a key consideration. Throughout this paper, I refer to the *optimal reserve size* in the sense of sustainable fishing yields. However, it should be noted that marine reserves can provide most fishery and conservation benefits at anywhere from short to long distances [35], including promotion of biodiversity [2, 45] and genetic diversity [7], improving ecological resilience [6, 52] and optimal fishery profit [47], mitigation of impacts from climatic change [21, 43] and catastrophic events [58], to name just a few.

Previous studies have revealed several important properties of the optimal reserve size. Using a model of population dynamics described by a stock-recruit relationship, Hart [25] determined that an optimal reserve size exists when the slope of the stock-recruit curve at a given fishing mortality rate is larger than the inverse of the harvest-free spawning stock biomass per recruit. Applying this principle to canary rockfish and Georges Bank sea scallop, Hart [25] showed that a fishing mortality rate greater than *f*_*MSY*_ is necessary to meet this condition. Takashina [49] adopted an age-structured metapopulation model and showed that an intermediate recruitment success of eggs and a large fishing mortality rate are required for marine reserves to improve fishing yields. Bensenane, Moussaoui, and Auger [8] analyzed a bioeconomic model that describes dynamics of population size and fishing mortality rate. They derived an algebraic formulation of an optimal reserve size, composed of the market price of the target species, catchability coefficient, the carrying capacity, and the cost of fishing effort, showing that an optimal reserve size is economically feasible. Furthermore, several studies [13, 17, 42, 57] showed numerically, in models where larval dispersal attributes the spillover from marine reserves, that implementation of marine reserves promote fishing yields compared to no-reserve management, supporting the existence of an optimal reserve size. However, previous studies concerning optimal reserve size have avoided the effect of adult movement [13, 25, 49] or have addressed only well-mixed populations [8, 57].

Adult movement significantly affects the performance of marine reserves via the spillover effect [22, 23, 35], including species abundance and fishing yields [14, 55], population persistence [31, 38], and spatial and temporal scales of population responses after reserve implementation [37]. Adult mobility increases population export from marine reserves, reducing conservation benefits and compensating for lost fishing opportunity [22]. Also, marine organisms show highly diverse mobility and dispersal behavior, including density-dependent and density-independent (diffusion) migrations [22]. For instance, fish and echinoderm species often exhibit density-dependent movement. This has been investigated experimentally [44] and through a field study [1]. Density-dependent movement heavily influences performance of marine reserves, such as fishery outcomes under various management scenarios (e.g., maximizing current profit, open access.) [4], ecological resilience [52], and population persistence [36]. Therefore, investigating the effect of adult movement on optimal reserve size, and understanding its relationship to conservation benefits, such as population sizes [19, 34], may further enhance fishery management success.

Here, I investigate the relationship between optimal reserve size and adult spillover, and realized conservation benefit, measured by the total population size. The model is a simple two-patch model, where one patch represents a fishing ground and the other, a marine reserve. Migration connects the two patches at a species-specific rate, and affects the degree of spillover from the marine reserve. This creates source-sink dynamics between the fishing ground and marine reserve, and this is a common structure of existing spatially-explicit models (e.g., [12, 18, 39, 52]). The two-patch model also allows us to examine different migration modes [3], such as positive/negative density-dependent, as well as density-independent migration. This simple approach yields analytical results by introducing an aggregated model when the migration rate is sufficiently larger than the growth rate of a target species [5,8,29,51,53]. The aggregated model allows us to derive conditions for optimal reserve size, fishing yield, and total population size under management with the optimal reserve fraction. I show that an optimal reserve size exists when the fishing mortality is larger than *f*_*MSY*_, and that the fishing yield and the total population size under the management correspond to MSY and *X*_*MSY*_, the population size at MSY.

In addition, I numerically examine the model with various migration rates and modes when model aggregation is not valid. In addition to fishing yields, I also calculate the total population size as a measure of the conservation benefit of marine reserves and discuss the tradeoff under management with marine reserves. I find that an optimal reserve size exists under various migration rates and modes. Optimal reserve size tends to increase with migration rate and approaches the optimal size of the aggregated model, where the optimal size is the largest.

## 2 Model

### 2.1 Basic model

The starting point is the commonly used Schaefer model [20, 48] in which population dynamics of the target species *x* are described with growth rate *r*, carrying capacity *K*, and fishing mortality rate *f* as follows:

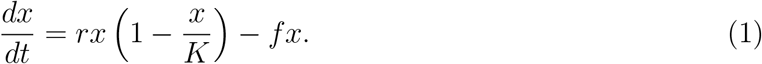

In this model, MSY, the fishing mortality rate at MSY, *f*_*MSY*_, and the population size at MSY, *X*_*MSY*_, are described as [11]

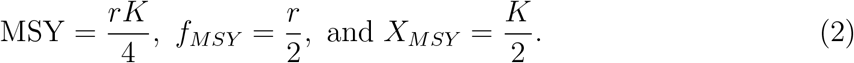

To investigate the effect of a marine reserve, I employ a common two-patch model (e.g., [18, 52, 54]) where one patch is open to fishing (*i* = 1; with fraction 1 − *α*), the other patch (*i* = 2; with fraction *α*) is protected from fishing (i.e., *f* = 0), and migration connects the two patches (Fig. 1). With the migration function *M* (*x*_1_, *x*_2_) where *x*_1_ and *x*_2_ are the population of the fishing ground and marine reserve, respectively, Eq. (1) becomes

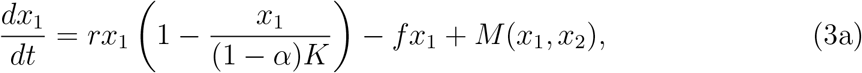

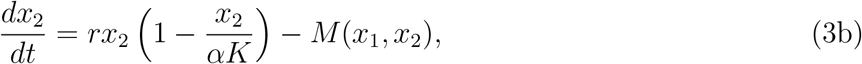

**Figure 1:**
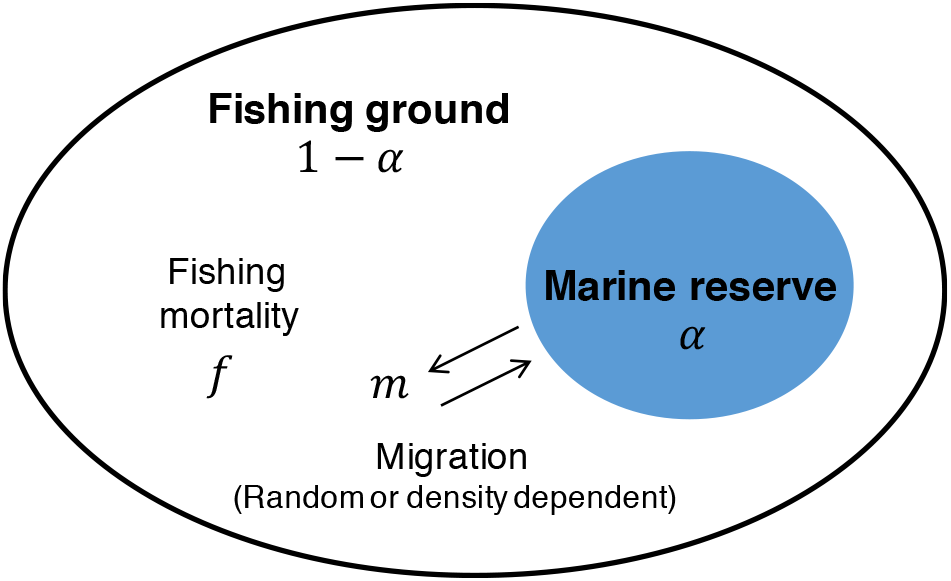
Schematic description of the model. The space is divided into the fishing ground (fraction 1 − *α*) with fishing mortality, *f*, and the marine reserve (fraction *α*) without fishing activity. Migration at rate *m* connects the dynamics of the two patches. Migration is either random or positively/negatively density-dependent.

I consider the situation where migrations between the patches are either random or positively/negatively density-dependent. With the migration rate between the marine reserve and the fishing ground, *m*, and the parameter controlling the migration mode, *s*, I describe the migration function *M* (*x*_1_, *x*_2_) as

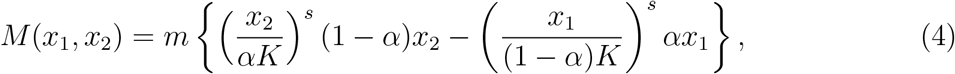

where, the function represents the negative density-dependent migration when −1 < *s* < 0, random migration when *s* = 0, and density-dependent migration when *s* > 0 [3].

I use the following notations to describe the fishing yield and the total population size under management with a marine reserve

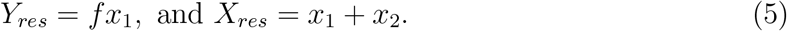

where, these quantities are used subsequently to compare management without a marine reserve.

### 2.2 Model aggregation

When the migration rate is sufficiently larger than the growth rate (*m* ≫ *r*), there are fast and slow dynamics operating at different time scales [5, 29]. Then, the migration term has a negligible effect on the total population *X* = *x*_1_ + *x*_2_ operated at the time scale of fast parameter *τ* = *mt*. Thus, Eq. (3) can be approximated by a single aggregated model [53]

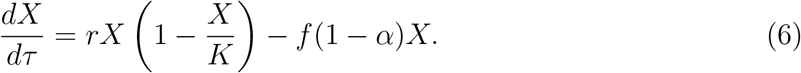

Hereafter, I discuss analytical aspects of the aggregated model (6), and perform numerical calculations across the migration rate *m*, including a situation where the model aggregation is not valid.

## 3 Results

### 3.1 Analysis of the aggregated model Eq. (6)

The aggregated model allows us to derive the explicit form of the equilibrium as follows:

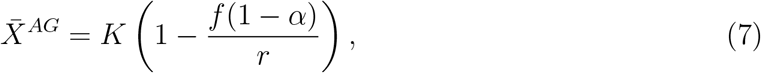

and fishing yield is

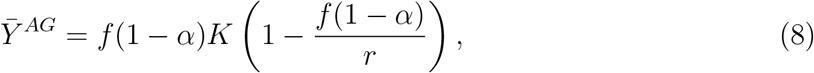

where, the superscript *AG* indicates the equilibrium of the aggregated model. By solving 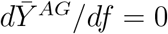 about *f* and *α*, respectively, I obtain the optimal fishing mortality and reserve size:

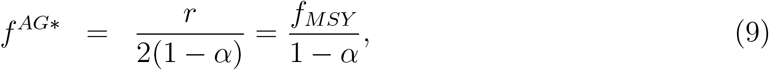

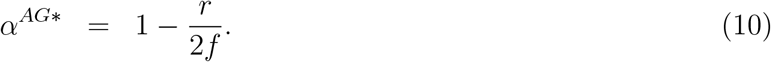

Eq. (9) suggests that one needs to increase the fishing mortality rate at the MSY level with a rate inversely proportional to the fraction of fishing ground 1 − *α* after establishment of a marine reserve. It becomes infinitely large when the fraction of the marine reserve approaches unity (Fig. 2). On the other hand, Eq. (10) shows that for an optimal reserve size to exist, the fishing mortality rate should be larger than MSY:

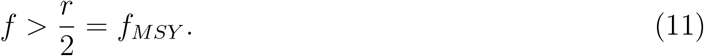

**Figure 2:**
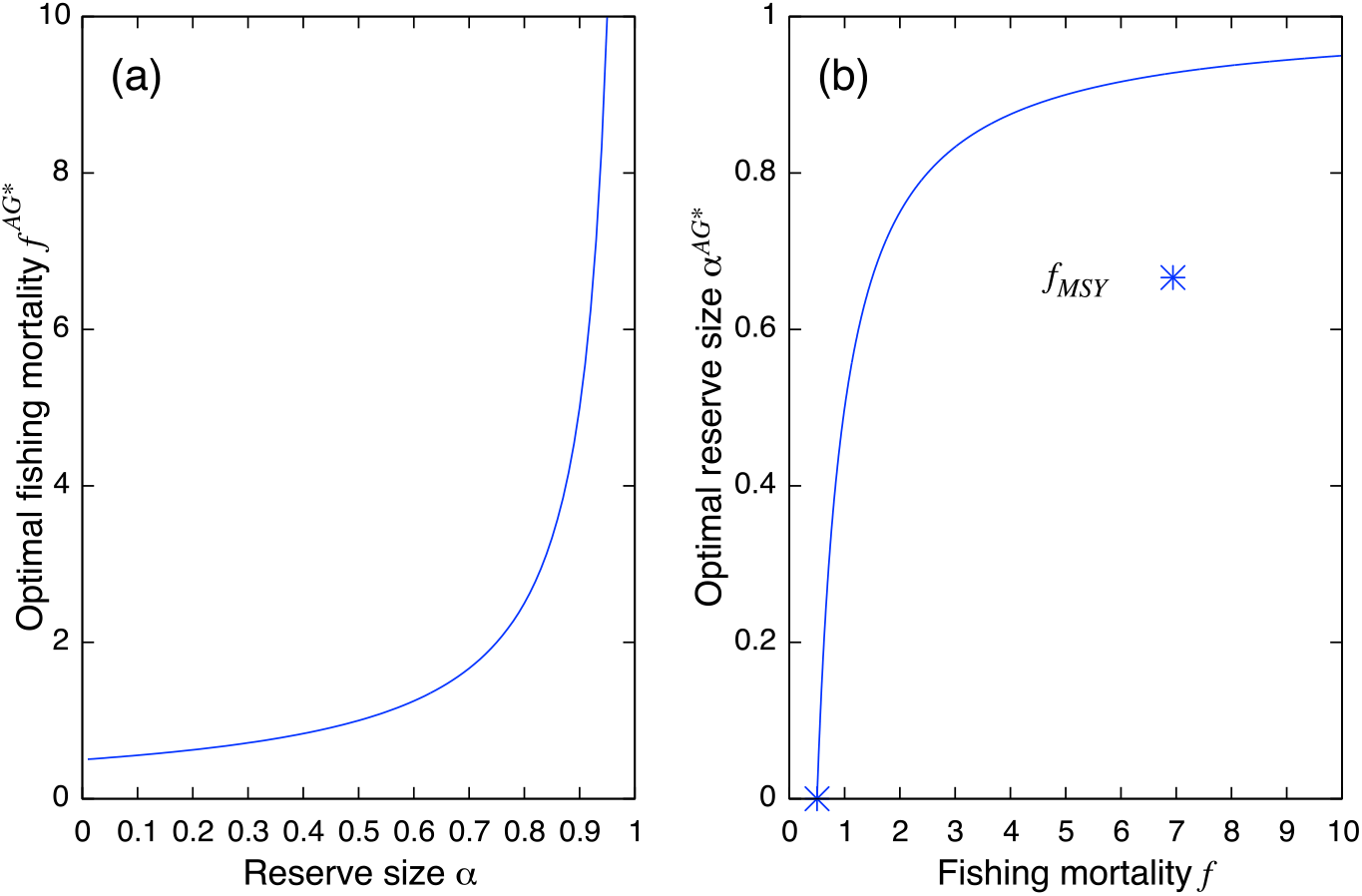
(a) Optimal fishing mortality of the aggregated model *f*^*AG*^* (Eq. (9)) and (b) optimal reserve size *α*^*AG*^* (Eq. (10)). Positive reserve size exists only when fishing mortality is larger than *f*_*MSY*_.

In other words, if fishing mortality is smaller than MSY, there is no optimal reserve size that can improve fishing yield. Also, the optimal reserve size approaches 1 (i.e., complete fishing ban) as fishing mortality becomes large (Fig. 2).

From Eq. (6), maximum fishing yields coincide with MSY of Eq. (1). In fact, substituting either Eq. (9) or (10) into Eq. (8) recovers Eq. (2). I denote this by

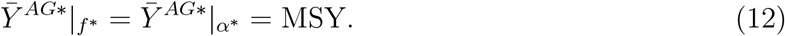

I often use notation 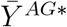 when the substitution is obvious. Similarly, I regard 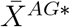 as the population size when fishing yield is given by Eq. (12).

### 3.2 Numerical investigation for general situation

When the migration rate is not large, then the aggregated model is not valid. Nonetheless, I will show that the analytical results above provide a maximum fishing yield, and this becomes a benchmark to discuss the performance of an introduced marine reserve.

Here, I numerically solve Eq. (3) to find the fishing yield and the optimal reserve size *α*\*, as well as the total population size under various reserve sizes and migration rates. To compare these quantities with those of management without a marine reserve, I introduce the following two normalized quantities: the fishing yield normalized by MSY, *Y*_*res*_/MSY, and the total population size normalized by *X*_*MSY*_, *X*_*res*_*/X*_*MSY*_. Note from the analysis above, the aggregated model under the optimal reserve size, *α*^*AG*^*, gives the values of normalized fishing yield and population size 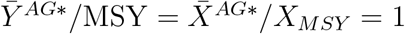.

The top three panels in Figure 3 show the normalized fishing yield under density-independent migration (*s* = 0). As expected, the optimal reserve size *α*\*, if any, approaches that of the aggregated model *α*^*AG*^* as the migration rate *m* becomes large. Also, the normalized fishing yield of the aggregated model gives the upper bound: 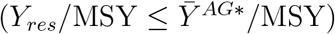. These calculations also suggest that the condition for the positive optimal reserve size exists (Eq. 11) still holds for various migration rates. Therefore, the improvement of fishing yield by introducing a marine reserve occurs only when the initial fishing mortality exceeds MSY, *f*_*MSY*_. Although this condition is necessary for a marine reserve to increase the harvest when the migration rate is not high enough (i.e., the model aggregation is not valid), it does not guarantee the existence of an optimal reserve size. For example, when fishing mortality is moderately high (*f* = 0.75; Figure 3b), a spillover effect from the marine reserve is necessary for an optimal reserve size to exist. A high fishing mortality rate (*f* = 1) relaxes this requirement (Figure 3c) and an optimal reserve size exists for all migration rates above *m* > 0.01. The bottom three panels in Fig. 3 show a slice of the heat map shown in the top panels at *m* = 0.1, 1, or 10. The fishing mortality rate *f* corresponds to the top panel. These demonstrate that the fishing yield tends to be higher when the migration rate *m* is high. Also, there are peaks when fishing mortality is larger than *f*_*MSY*_ = 0.5, and these correspond to the optimal reserve size *α*\*.

**Figure 3:**
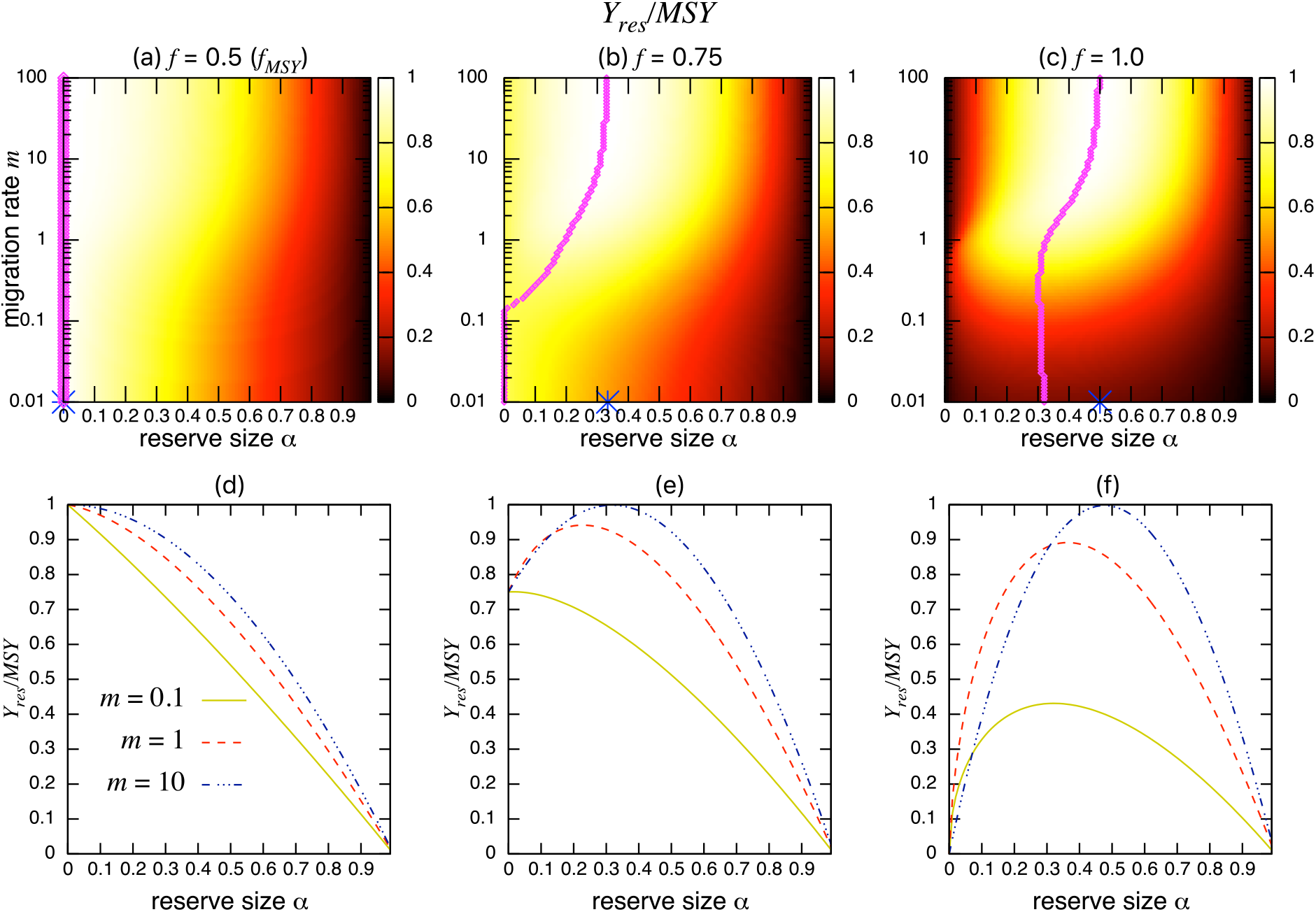
Effect of a marine reserve on fishing yield across reserve sizes *α* and migration rates *m* when migration is density-independent. (Top) the normalized fishing yield (*Y*_*res*_/MSY) and optimal reserve size *α*\* (magenta points (color online)). Lighter heat map color indicates a larger value in *Y*_*res*_/MSY. Optimal reserve size predicted by the aggregated model Eq. (6), *α*^*AG*^*, is also shown on *x*-axis (blue star (color online)). (Bottom) Slices of the heat map above at *m* = 0.1, 1, and 10. Fishing mortality *f* corresponds to the value shown in the top panel. Parameter values used are *r* = 1, *K* = 10, and *s* = 0.

On the other hand, the bottom three panels of Figure 4 show the normalized total population size. This value represents the conservation benefit of the marine reserve. Management with an optimal marine reserve size tends to give a smaller population size than *X*_*MSY*_ (i.e., *X*_*res*_*/X*_*MSY*_ < 1). However, if fishing mortality is larger than *f*_*MSY*_, this value approaches 1, and the prediction of the aggregated model, as the migration rate *m* becomes sufficiently large (Fig. 4b, c). Also, increasing the reserve size provides a higher normalized population size, and the population size becomes larger at low to moderate migration rates (*m* is about 1 in Figure 4) at a given reserve size. The bottom three panels in Fig. 4 show more explicitly this relationship when the migration rate is *m* = 0.1, 1, or 10. Unlike fishing yields, the population size, a measure of conservation benefit, is larger when the migration rate is moderate (*m* = 1) rather than high (*m* = 10). This suggests a mismatch between optimal harvesting and the conservation benefit.

**Figure 4:**
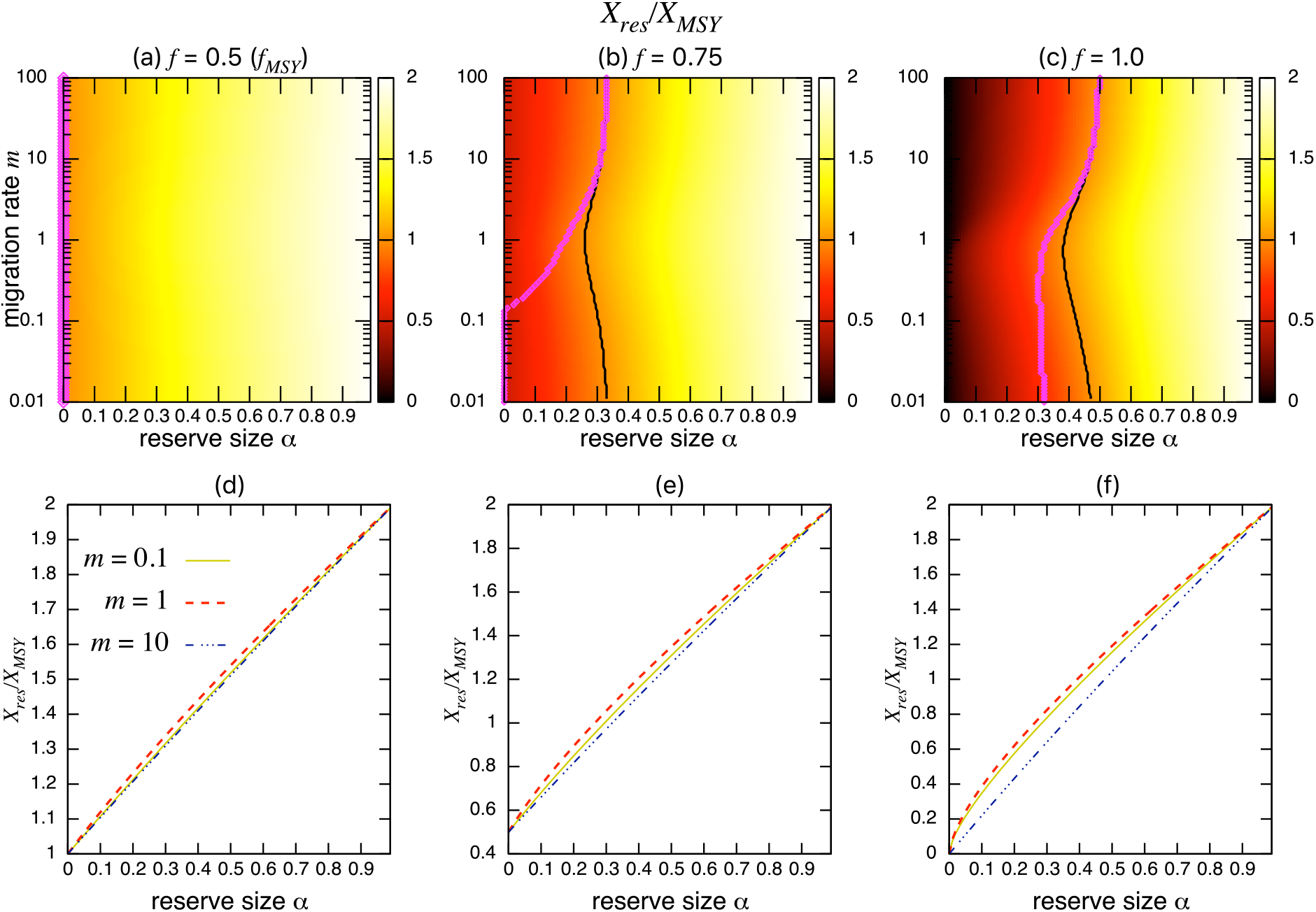
Effect of a marine reserve on population size at various reserve sizes *α* and migration rates *m* when the species migration is density-independent. (Top) Normalized population size (*X*_*res*_*/X*_*MSY*_) and optimal reserve size *α*\* (magenta points (color online)). Lighter color in the heat map indicates larger values in *X*_*res*_*/X*_*MSY*_. The black line represents the point where *X*_*res*_*/X*_*MSY*_ = 1. (Bottom) Slices of the heat map above at *m* = 0.1, 1, and 10. Fishing mortality *f* corresponds to the value shown the top panel. Parameter values used are the same as in Fig. 3.

I found qualitatively similar trends in two alternative migration modes: density-dependent (*s* = 1; Fig. 5) and negatively density-dependent (*s* = −0.5; Fig. 6) migrations between patches where I show only the heat maps. Some quantitative differences appear when the migration rate *m* is around 1, but these approach the prediction of the aggregated model as the migration rate becomes suffciently large, and optimal reserve size appear when fishing mortality is larger than *f*_*MSY*_.

**Figure 5:**
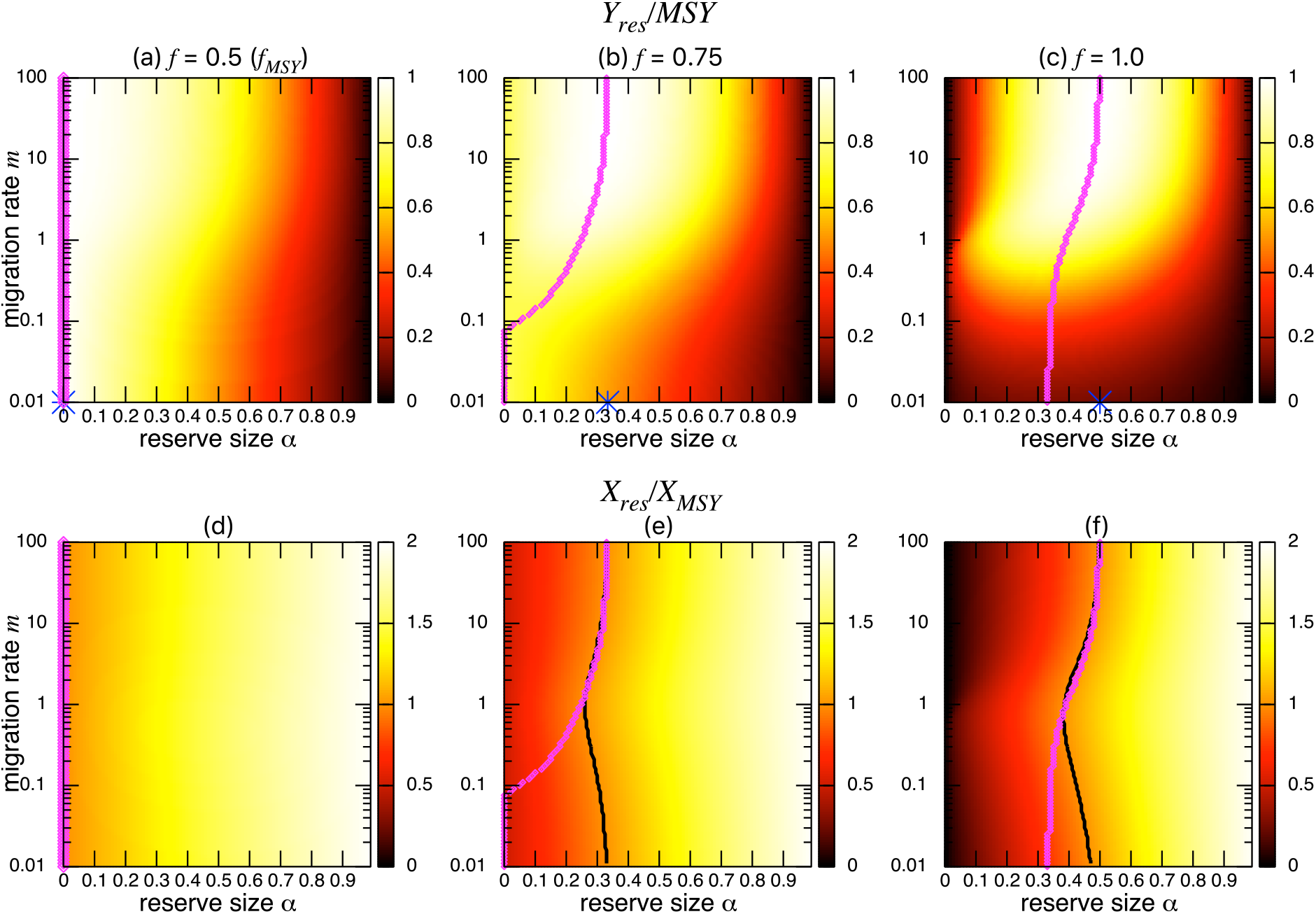
Effect of a marine reserve on the fishing yield and population size at various reserve sizes *α* and migration rates *m* when the migration is positively density-dependent (*s* = 1). (Top) the normalized fishing yield (*Y*_*res*_/MSY). Optimal reserve size predicted by the aggregated model Eq. (6), *α*^*AG*^*, is also shown on *x*-axis (blue star (color online)). (Bottom) the normalized population size (*X*_*res*_*/X*_*MSY*_). The black line represents the point where *X*_*res*_*/X*_*MSY*_ = 1, and the magenta points (color online) show the optimal reserve size *α*\*. Other parameter values used are the same as in Fig. 3.

**Figure 6:**
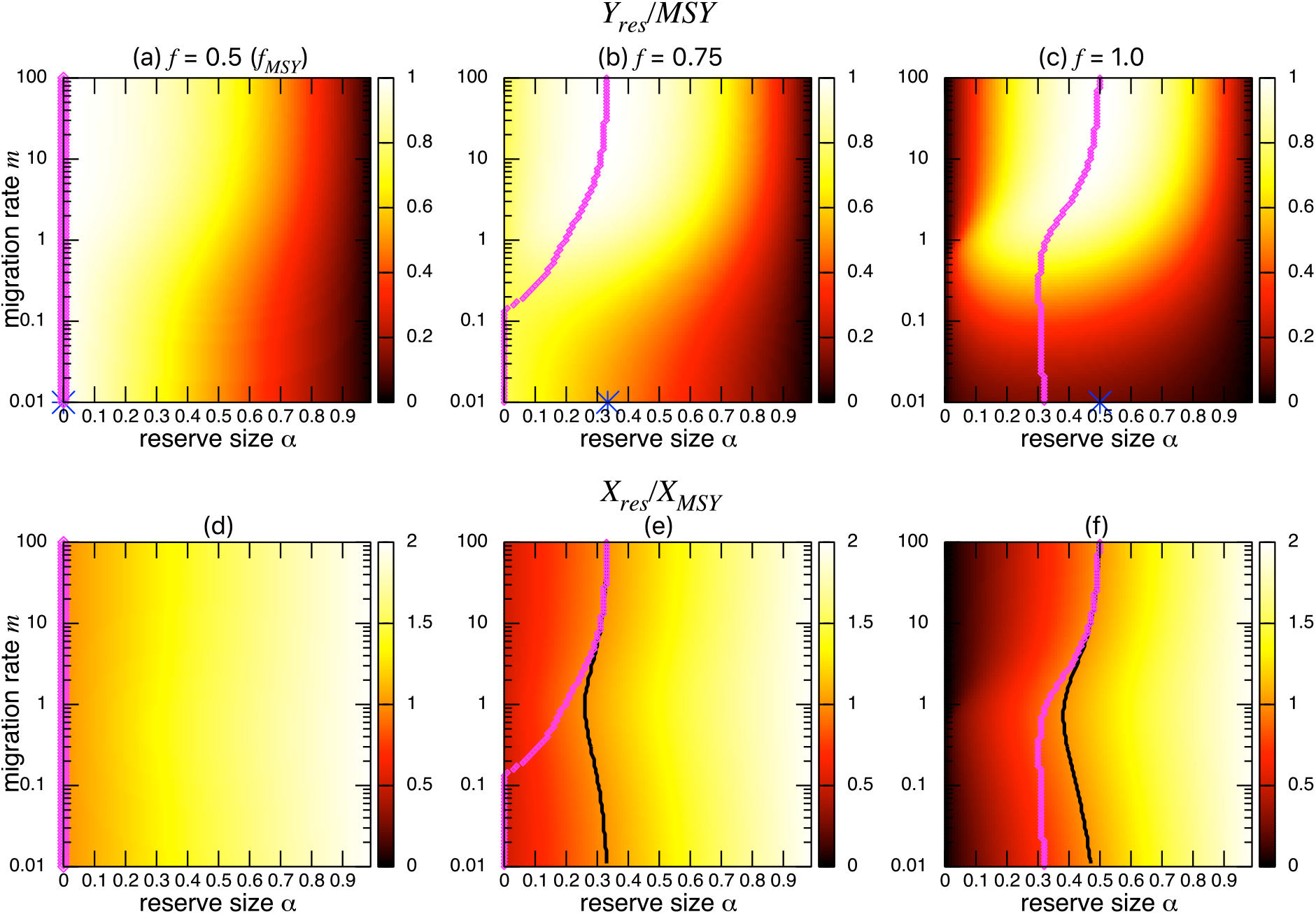
Effect of a marine reserve on fishing yield and population size at various reserve sizes *α* and migration rates *m* when the species migration is negatively density dependent (*s* = −0.5). (Top) the normalized fishing yield (*Y*_*res*_/MSY). Optimal reserve size predicted by the aggregated model Eq. (6), *α*^*AG*^*, is also shown on *x*-axis (blue star (color online)). (Bottom) the normalized population size (*X*_*res*_*/X*_*MSY*_). The black line represents the point where *X*_*res*_*/X*_*MSY*_ = 1, and the magenta points (color online) show the optimal reserve size *α*\*. Other parameter values used are the same as in Fig. 3.

## 4 Discussion

The effect of a marine reserve on fish catch is a crucial consideration because creation of a marine reserve in an existing fishing ground initially reduces fishing opportunities. By taking advantage of a simple two-patch model, I derived a necessary condition for an optimal size of the marine reserve to exist (*f* > *f*_*MSY*_; Eq. (11)) confirming a previously reported value [25, 40, 42]. I also derived a simple formula for optimal fishing mortality rate (Eq. (9)) and optimal reserve size (Eq. (10)). Numerical simulations examined various migration rates and modes and revealed several new findings. That is, the theoretically predicted fishing yield from the aggregated model gives an upper bound: a well-mixed population with a large migration rate maximizes the fishing yield. In addition, even with the condition *f* > *f*_*MSY*_, a moderate migration rate is necessary, on top of previous findings [25, 40, 42], for an optimal reserve size to exist when fishing mortality is moderately high (Fig. 3b), but this additional condition is not necessary when fishing mortality is high (Fig. 3c). I also found that different migration modes lead to qualitatively similar results, a high migration rate decreases the effect of different migration modes, and fishing yields and population sizes approach values predicted by the aggregated model. On the other hand, the model shows that a marine reserve provides a larger total population, a measure of conservation benefit when the migration ability of a target species is low or moderate, contrasting to benefits for fishing yield. This emphasizes the importance of spillover effects in considering the tradeoffs under management of marine reserves.

The necessary condition derived from the aggregated model for marine reserves to improve fishing yields confirms a condition previously reported [25, 40, 42] under different modeling frameworks, suggesting potentially generic characteristics of marine reserve management. If a marine reserve replaces a certain fraction of a fishing ground, the fishing mortality rate at the MSY level should increase by a factor inversely proportional to the fraction of the fishing ground, 1 − *α*. In practice, this corresponds to a situation where all fishermen remain in the contracted fishing ground, with proportion 1 − *α*, after reserve creation. Under these conditions, the fishing yield becomes as large as MSY [8, 26]. These results suggest that the optimal reserve size spans *α*\* ∈ (0, 1), and the optimal reserve size approaches 1 as fishing mortality becomes sufficiently large. In practice, however, excluding most fishing from a region of concern may be infeasible with multiple management tradeoffs. In fact, the often suggested range of optimal reserve size to increase fishing yields and/or to meet management and conservation objectives is smaller; e.g., about 30% of the region of concern in the synthetic analysis [16] or 20% − 50% in [24]. My results suggest that the maximum harvest under management with a marine reserve is achieved when the species migration rate becomes sufficiently large (*m* ≫ 1). This indicates that the reduced fishing ground can be compensated by increased fishing mortality when there is sufficient migration from a marine reserve to a fishing ground. However, intensified fishing mortality to achieve the maximum fishing yield given a migration rate *m* often leads to a smaller total population than the MSY: *X*_*res*_*/X*_*MSY*_ < 1, except for the case *m* ≫ 1 where *X*_*res*_*/X*_*MSY*_ = 1. On the other hand, the model shows a larger population size at an intermediate migration rate (Fig. 4b and c), representing the underlying tradeoff between fishing yield and population size (i.e., conservation benefit). Previous studies also showed that marine reserves may provide larger conservation benefits, such as a larger population recovery and reproductive capacity, at relatively low to moderate migration rates [9, 22, 52].

The finding that high migration rates (*m* ≫ 1) produce the highest fishing yields implies a close relevance to the results of Neubert [39], where the number of marine reserves to maximize the fishing yield can become infinitely many in a finite length of one-dimensional space, leading to an infinitely large exchange rate between marine reserves and fishing grounds. It suggests that in practice, migration between marine reserves and non-reserve sites can be controlled by the configuration of marine reserves in space, and that the migration rate is not a purely biological parameter. Therefore, there may exist a marine reserve design to realize a high migration rate in the setting of my model, even for a sedentary species by, for example, creating smaller marine reserves than the home range sizes of target species. Although the two-patch model simplifies the marine environment, and the migration rate *m* is constant regardless of the reserve size *α*, investigation of the influence of migration ability (e.g., diffusivity) and mode of a target on the migration rate, *m*, will further improve marine reserve management. The two-patch model, however, is still a useful framework to discuss the performance of marine reserves with a simple function of migration terms, enabling us to avoid the description of complex migratory dynamics within each patch. Rosenberg et al. [44] experimentally demonstrated that a brittle star generally shows density-dependent dispersal and estimated dispersal speed. Therefore, it may be possible to perform an experiment to estimate the parameter controlling density dependence. Also, the fact that the configuration of marine reserves influences the migration rate highlights the importance of considering the spatial scaling of management. Previously, Takashina and Baskett [50] demonstrated that the management unit scale is decomposed from the spatial scale in which biological processes operate. They showed that fisheries management with a finer management unit scale results in larger fishing yields than management with a larger management scale, since the former can realize a fine-tuned allocation of fishing efforts across management units. Fine-tuned fishing activities across fishing grounds allows higher flexibility in management decision making than the two-patch model, where I can assign only a single fishing mortality rate. Hence, consideration of the migration rate as a function of the management unit scale may offer a more flexible way to mitigate underlying management tradeoffs.

Prediction of the equivalence in MSY and the yield from management involving marine reserves has been revised by preceding studies showing that marine reserve result in a fishing yield larger than MSY. For example, Gaylord et al. [17] and Ralston and O’Farrell [42] attributed excess yields to spatial structures of the model and post-dispersal density-dependent recruitment of larvae, and De Leo and Micheli [13] demonstrated that a highly complex spatially-explicit, stage-structured stochastic model improves fishing yields. White and Kendall [57] claimed that model complexity is not responsible for this result, and they demonstrated that only the effect of post-dispersal, density-dependent recruitment in population dynamics induces excess fishing yields. On the other hand, my simple mathematical model does not show a fishing yield larger than MSY. The lack of detailed spatial structures, stage/age-structure, and/or post-dispersal density-dependent recruitment, as in the above examples, may explain this contrasting result. Particularly, species migration and larval dispersal cause a rather different population mixing pattern. For example, post-dispersal, density-dependent recruitment causes a spontaneous reduction of the number of larvae recruited, and it is often assumed to occur at a discrete-time (i.e., seasonal breeding cycle) [13, 17, 42, 57], contrasting with my continuous-time model where population exchanges and density-dependent population control occur throughout the year. In marine environments connected by larval dispersal, less mobile species are more likely to increase fishing yields than highly mobile species. That is, sessile species in reserves tend to stay in the reserve longer; hence, they receive greater benefits from reserves and provide more recruits to fishing grounds [28, 59]. On the other hand, high adult migration rate causes frequent population exchanges between reserves and fishing grounds, and such species are more vulnerable to fishing activities than those with lower migration rates, leading to higher fishing yields than for less mobile taxa. Provided that the optimal yield in the aggregated model is the upper bound in the two-patch model, the model serves as a basic model to assess the effect of stage/age-structure, density-dependent larval recruitment, and more complex fishing regulations under different levels of spillover. For example, size/age-based regulation is prevalent in fisheries management, and its explicit consideration with various species migration scenarios further improves our understanding of optimal reserve size.

In this paper, I used the simplest possible model to discuss optimal reserve size while accommodating various adult migration rates and modes. Optimal yields provide a useful benchmark to assess the strength of management tradeoffs. There are, of course, a number of different mathematical approaches possible (See the review by [15] for models used in studies of marine reserves). With a good understanding of adult movement of a species of concern, one natural extension of the two-patch model is the integration of metapopulation structure, as discussed in [31, 32] that realizes more complex pathways of species and enables more realistic conservation planning. Multiple-species dynamics are also important components of marine ecosystem management [41,54] and discussion of the impact of species interactions on optimal reserve sizes will improve our insight. Understanding these effects on optimal reserve management, together with adult movement, will be necessary.

## Acknowledgements

NT was financially supported by Grant-in-Aid for the Japan Society for the Promotion of Science Fellows. I thank reviewers for their thoughtful comments.

